# Sex classification by resting state brain connectivity

**DOI:** 10.1101/627711

**Authors:** Susanne Weis, Kaustubh Patil, Felix Hoffstaedter, Alessandra Nostro, B.T. Thomas Yeo, Simon B. Eickhoff

**Author notes:** Corresponding author: Susanne Weis, Institute of Systems Neuroscience, Medical Faculty, Heinrich Heine University Düsseldorf, Düsseldorf, Germany; Institute of Neuroscience and Medicine, Brain & Behaviour (INM-7), Research Centre Jülich, Jülich, Germany.

## Abstract

A large amount of brain imaging research has focused on group studies delineating differences between males and females with respect to both cognitive performance as well as structural and functional brain organization. To supplement existing findings, the present study employed a machine learning approach to assess how accurately participants’ sex can be classified based on spatially specific resting state (RS) brain-connectivity, using two samples from the Human Connectome Project (n1 = 434, n2 = 310) and one fully independent sample from the 1000BRAINS study (n=941). The classifier, which was trained on one sample and tested on the other two, was able to reliably classify sex, both within sample and across independent samples, differing both with respect to imaging parameters and sample characteristics. Brain regions displaying highest sex classification accuracies were mainly located along the cingulate cortex, medial and lateral frontal cortex, temporo-parietal regions, insula and precuneus. These areas were stable across samples and match well with previously described sex differences in functional brain organization. While our data show a clear link between sex and regionally specific brain connectivity, they do not support a clear-cut dimorphism in functional brain organization that is driven by sex alone.

## 3 Introduction

A large amount of brain imaging research has focussed on delineating differences between males and females with respect to both cognitive performance as well as structural and functional brain organization. However, while the terms “male brain” and “female brain” are often used both in scientific and popular writing, it is so far unclear if a sexual dimorphism in the human brain actually exists. In a strict sense the term “dimorphism” should only be used for those aspects of differences that come in two strictly distinct forms like the male and female genitalia (Joel D and A Fausto-Sterling 2016). In contrast, it has been suggested that most differences in brain and behaviour are not actually dimorphic, since they show a high degree of overlap between males and females (Joel D and A Fausto-Sterling 2016). With respect to brain structure, some literature (Joel D et al. 2015) even argues that any particular brain might comprise certain features that are statistically more typical of females and others which are more typical for males. In that sense, these authors suggested (Joel D *et al.* 2015), that most brains are comprised of “mosaics” of features, some more common in females, some more common in males, and some common in both. They showed that brains with structural features that are consistently at either end of the “maleness-femaleness” continuum are rare and an extensive overlap exists between the distributions of females and males for grey matter, white matter, and structural connectivity (Joel D *et al.* 2015). However, several other authors have criticized this conclusion by showing that multivariate classifiers can be trained to classify male and female brains based on structural data with a high accuracy of roughly 80% (Chekroud AM et al. 2016; Del Giudice M et al. 2016; Rosenblatt JD 2016). These data suggest that despite the absence of dimorphic differences and lack of internal consistency observed by (Joel D *et al.* 2015), multivariate analyses of whole-brain structural patterns are able to reliably classify the sex of a subject. Of note, these two results are not mutually inconsistent (Chekroud AM *et al.* 2016). While a strict dichotomy between the brains of males and females might not exist, this does not mean that statistical differences cannot or should not be considered (Chekroud AM *et al.* 2016).

Recent group studies also reported structural differences between the sexes. For example, the so-far largest single-sample study on sex differences in the brain, comprising more than 5200 participants (Ritchie SJ et al. 2018), found that males had higher cortical and sub-cortical volumes, cortical surface areas and white matter diffusion directionality while females had thicker cortices and higher white matter tract complexity. Furthermore, these authors identified some subregional differences that were not fully attributable to differences in total volume, total surface area, mean cortical thickness, or height. Similarly, a meta-analysis of more than 100 studies (Ruigrok AN et al. 2014) showed that, on average, males have larger total brain volumes than females and identified regional sex differences in volume and tissue density in the amygdala, hippocampus and insula.

Despite these structural differences, it is unclear, if and how structural brain variations translate into differences in functional brain organization. While a large body of literatures has aimed to delineate the cognitive domains in which males and females differ, assessment of brain function using functional imaging techniques has so far mostly revealed relatively small and inconstant group-level differences between females and males. Historically, sex differences have been reported across a wide variety of cognitive tasks (Miller DI and DF Halpern 2014) and a variety of brain imaging studies have been conducted to identify the brain basis of these difference. Implicitly assuming that there actually is a binary distinction between the male and female brain, the vast majority of these studies are based on group comparisons between males and females. However, results so far have been inconclusive (Hyde JS and EA Plant 1995; Del Giudice M 2009). For certain cognitive domains, especially language and emotional processing, there has been some evidence of sex differences in cognition and functional brain organization. However, more recent, larger analyses have not found any conclusive sex effects in these domains (Russell TA et al. 2007; Wallentin M 2009).

Further trying to elucidate the neuronal basis of sex differences, more recent functional magnetic resonance imaging (fMRI) research has considered functional brain connectivity in the absence of any specific cognitive task (Weis S et al. 2017; Ritchie SJ *et al.* 2018; Zhang C et al. 2018). Resting state (RS) fMRI provide an estimate of the functional connectivity of the brain at rest, i.e. the intrinsic brain connectivity. Some studies have identified specific networks, in which RS connectivity seems to differ between the sexes (Biswal BB et al. 2010; Zuo XN et al. 2010; Tian L et al. 2011; Ritchie SJ *et al.* 2018). However, other studies have not found any effect of sex in RS fMRI data (Weissman-Fogel I et al. 2010). Furthermore, a more recent study, employing repeated RS measurement across different menstrual cycle phases in women and time-matched tests in men, identified sex differences in some RS networks, while in others, the sex difference was dependent on the cycle phase of the women (Weis S *et al.* 2017), suggesting that sex is not the only factor influencing individual differences in RS connectivity.

### 3.1 Sex Classification

Altogether, evidence for sex differences in functional brain organization is inconsistent. Importantly, existing results based on group studies are fundamentally compromised by being based on the questionable assumption of a clear-cut sexual dimorphism of the human brain. Thus, more advanced computational methods seem to be more appropriate for the characterization of the complex patterns that characterize differences between the sexes. To this end, machine-learning methods can be used to delineate how accurately the sex of an individual, out-of-sample subject can be predicted from neuroimaging data (Bzdok D 2017). In this approach, a classifier learns the relationship between a set of features, which are extracted from brain imaging data, and a particular outcome, in this case the sex of the subject, using a sample of observations. In the next step, this classifier can be used to predict the sex of a previously unseen subject given its features.

To date, there are not many studies that have adopted a classification approach based on structural (Feis DL et al. 2013; Chekroud AM *et al.* 2016; Rosenblatt JD 2016) or functional (Smith SM, D Vidaurre, et al. 2013; Ktena SI et al. 2018; Zhang C *et al.* 2018) brain imaging data. In general, these studies employed whole brain structural and or functional connectivity based on pre-defined regions of interest (ROIs) or brain parcellations. Based on whole-brain connectivity patterns, they have achieved a sex prediction accuracy of roughly 80% both based on brain structure and function. While these findings suggest that multivariate analyses of whole-brain structural patterns are able to reliably classify the sex of a subject based on brain imaging data, whole brain functional connectivity does not appear to be the optimal approach to characterize the brain basis of sex differences.

From a methodological point of view, machine-learning approaches based on whole brain connectivity are extremely vulnerable to the so called “curse of dimensionality”. Features based on whole-brain connectivity are of extremely high dimensionality. Even if the connectivity patterns are computed based on a parcellation of only a few hundred parcels, the dimension of connectivity feature vectors ranges on the order of several tens of thousands – much higher than the dimensionality of the data sets which usually contain far less than a thousand subjects. To avoid overfitting resulting in deflated classification accuracies, dimensionality reductions need to be applied. Furthermore, whole brain connectivity complicates the interpretability of the results, as it is difficult to conclude which specific parts of the brain are most distinct between males and females. As classification performance can usually not be linked to specific brain regions, it is impossible to draw any conclusions as to those cognitive domains in which males and females differ most. Thus, it is impossible to put sex classification findings in relation to findings from classical group studies.

To avoid the curse of dimensionality, while at the same time aiming to identify spatially specific effects, we employed a novel approach that is based on spatially specific connectivity for individual ROIs across the brain, instead of being based on whole brain connectivity. The examination of spatially specific effects is based on the assumption that sex differences in the performance within specific cognitive domains can be taken to suggest some rather selective neural differences restricted to specific brain regions. As opposed to most previous studies, which employed on whole brain connectivity for classification, we chose a parcelwise approach to assess how many and which brain regions’ connectivity is best able to classify sex with highest accuracies. Identifying those brain parcels that achieve high accuracies based on their connectivity patterns allows for straight-forward interpretations, especially when putting the present results in relation to classical group studies on sex differences. Furthermore, our parcelwise approach avoids the curse of dimensionality, which typically leads to worsened predictions in very high dimensional data sets like the whole brain connectome.

Altogether, the present study aimed to show that parcelwise connectivity patterns allow the classification of previously unseen subjects sex with accuracies that approach those that can be achieved based on whole brain connectivity. Based on spatially specific effects, functional decoding, i.e. meta-analyses based analysis of structure-function relationships (Fox PT et al. 2014), can be employed to identify the cognitive domains that these brain regions are related with. Only the assessment of such spatially specific effects makes it possible to directly link sex classification results to sex differences in specific cognitive domains as suggested by existing group studies.

Additionally, we aimed to examine if these spatially specific effects generalize across samples, differing both with respect to imaging parameters and sample characteristics like age. To this end, we trained a classifier on one set of imaging data and applied this classifier to an independent data set. If the classifier performs well on independent samples, this can be taken as strong support for the generalizability of spatially specific brain differences between the sexes.

## 4 Materials and Methods

### 4.1 Samples

Two mutually exclusive samples of unrelated subjects were constructed from data provided by the Human Connectome Project (HCP S1200 release, Van Essen et al., 2012). Sample 1 contained 434 subjects (age range: 22-37, mean age: 28.6 years, 217 males), sample 2 comprised 310 subjects (age range: 22-36, mean age: 28.5 years, 155 males). Within each of the two samples, males and females were matched for age, twin-status and education. Twins were not included in the samples. Resting state (RS) BOLD data comprised 1200 functional volumes per subject, acquired on a Siemens Skyra 3T scanner with the following parameters: voxel size= 2 × 2 × 2 mm³, FoV= 208 × 180 mm², matrix = 104 × 90, 72 slices, TR = 720 ms; TE= 33.1 ms, flip angle =52°. The data were collected using a novel multi-band echo planar imaging pulse sequence that allows for the simultaneous acquisition of multiple slices (Xu J et al. 2013). For RS data acquisition, subjects were asked to lie with eyes open, with “relaxed” fixation on a white cross (on a dark background), think of nothing in particular, and not to fall asleep (Smith SM, CF Beckmann, et al. 2013). Sample 3, a fully independent sample covering a different age range, was obtained from the population-based 1000BRAINS study (Caspers S et al. 2014). It comprised 300 volumes per subject, scanned on a Siemens TRIO 3T scanner with the following parameters: voxel size= 3.1 × 3.1 × 3.1 mm³, FoV= 200 × 200 mm², matrix = 64 × 64, 36 slices, TR = 2200 ms; TE= 30 ms, flip angle =90°. This sample comprised 941 subjects (age range: 18 – 88, mean age: 62.8 years, 512 males). During RS data acquisition, participants kept their eyes closed and were instructed to let the mind wander without thinking of anything in particular (Caspers S *et al.* 2014).

To examine the influence of volumetric differences between males and females, an additional sample (sample 4) was created from sample 1 and 2 (both HCP samples), in which males and females were matched for grey matter volumes. This new sample comprised 260 participants (age range: 22 – 37, mean age: 28,48, 130 males).

### 4.2 Pre-processing

For sample 1 and sample 2, we employed the pre-processed and FIX-denoised data provided by the Human Connectome Project (HCP S1200 release), for which also the spatial normalization to the MNI152 template had already been performed before download. Thus, no further motion correction was performed. Movement parameters, as provided in the HCP S12000 release, indicated that movement in the scanner, measured as mean framewise displacement (FD, (Power JD et al. 2014)) did not differ between males and females in sample 1 (females: mean(FD) = 0.164, SD(FD) = 0.063, males: mean(FD) = 0.171, SD(FD) = 0.073, t = 1.038, p >0.05) or sample 2 (females: mean(FD) = 0.166, SD(FD) = 0.053, males: mean(FD) = 0.157, SD(FD) = 0.055, t = 1.364, p >0.05).

For sample 3, to ensure that physical noise and effects of within scanner motion are minimized as much as possible, RS fMRI data were cleaned of structured noise through the Multivariate Exploratory Linear Optimized Decomposition into Independent Components (MELODIC) method from the FSL toolbox (www.fmrib.ox.ac.uk/fsl). This process combines independent component analysis with a more complex automated component classifier referred to as FIX (FMRIB’s ICA-based X-noisifier) to automatically remove artefactual components (Salimi-Khorshidi G et al. 2014). The FIX-denoised data were further pre-processed using SPM12 (Statistical Parametric Mapping, Wellcome Department of Imaging Neuroscience, London, UK, http://www.fil.ion.ucl.ac.uk/spm/), running under Matlab R2014a (Mathworks, Natick, MA). For each participant, the first four echo-planar imaging (EPI) volumes were discarded prior to further analyses. Then EPI images were corrected for head movement by affine registration using a two-pass procedure: in the first step, images were aligned to the first image, and in the second step to the mean of all volumes.

Movement in the scanner, measured as mean framewise displacement (FD, (Power JD *et al.* 2014)) did not differ between males and females (females: mean(FD) = 0.132, SD(FD) = 0.001, males: mean(FD) = 0.130, SD(FD) = 0.001, t = 0.321, p >0.05). The mean EPI image was spatially normalized to the MNI152 template (Holmes CJ et al. 1998) by using the “unified segmentation” approach in order to account for inter-individual differences in brain morphology (Ashburner J and KJ Friston 2005). This approach was chosen, as several recent studies have indicated increased registration accuracies of this approach as opposed to normalization based on T1 weighted images (Calhoun VD et al. 2017; Dohmatob E et al. 2018).

### 4.3 Connectome Extraction

Instead of using whole brain connectivity, as previous studies have done, our novel approach is based on training classifiers on individual brain regions’ connectivity with the rest of the parcels. This approach allows the computation of sex classification accuracies individually for each of the regions to find out which brain areas’ connectivity achieve the best classification accuracies.

For each parcel, an activation time course was computed and correlated with those of each of the other parcels. Then, for each parcel individually, the connectivity pattern with the rest of the brain was used as features to train a classifier to distinguish between males and females. Finally, by using cross-validation, for each parcel the out-of-sample accuracy of sex classification was determined. This novel approach thus offers a straightforward way to delineate spatially specific effects.

Individual RS connectomes were created based on a novel whole-cortex parcellation reported by (Schaefer A et al. 2017), comprising 400 parcels. Since this atlas does not cover subcortical structures, we added 36 subcortical parcels taken from the Brainnetome atlas (Fan L et al. 2016). The Schaefer parcels have been shown to agree with the boundaries of certain cortical areas defined using histology and visuotopic fMRI, revealing neurobiologically meaningful features of brain organization. The Brainnetome atlas is fine-grained and cross-validated containing information on both anatomical and functional connections. The time-series of each parcel was cleaned by excluding variance that could be explained by mean white matter and cerebrospinal-fluid signal (Satterthwaite TD et al. 2013). For each parcel, the subject-specific time-series was then computed as the first eigenvariate of the activity time courses of all voxels within the parcel. For each parcel, we then computed pairwise Pearson correlations between the parcel’s time series with those of all other parcels, which were then transformed to Fischer’s Z-scores. Each parcel’s connectivity with the 435 other parcels across the whole brain for each subject was used as features in the classification analysis for this specific parcel.

### 4.4 Sex Prediction

For each brain parcel individually, nonlinear SVM (LibSVM toolbox, (Chang CCL, C.L. 2011)) with RBF kernel was employed to train a model for classification of the subject’s sex from the corresponding connectome. SVM learns the relationship between a set of input variables or features (the connectivity pattern of each individual parcel), and a particular outcome (the sex of the subject) across a set of observations. Our goal here was to fit a function that approximates the relation between the features and the outcomes, which can be used later on to infer the sex of new subjects from their connectome. Effects of age were adjusted using betas fitted only in the training. In an inner loop, the hyper-parameters gamma and C of the model were optimized by employing a cross-validation on the training set for each fold and the final model was created by averaging the hyperparameters across folds. For each parcel individually, we trained the classifier on sample 1 and determined within sample accuracy by a 10-fold cross validation, where the classifier was trained on 90% on the sample and tested on the remaining 10%. The same analysis was conducted for sample 4.

To characterize the statistical significance of the results an approximate permutation test approach was employed, in which associations between features and labels (sex) were randomized. That is, the labels were randomly permuted while the feature matrix was kept unchanged. 10-fold cross validation was repeated for each permutation and accuracies for 5000 permutations were used to construct an empirical null distribution. To control for multiple comparisons across the 436 parcels, the maximum accuracies across all brain parcels were obtained, resulting in 5000 null values obtained from the 5000 permutations, which were then used to compute FWE corrected p-values.

In the second step, the classifier was trained on the full sample 1 and then tested on sample 2 and sample 3. The brain networks were visualized with the BrainNet Viewer (http://www.nitrc.org/projects/bnv/) (Xia M et al. 2013).

### 4.5 Functional decoding

By training independent SVMs or each parcel of the brain, we are able to identify, to what extend connectivity of each parcel differentiates between males and females. These spatially specific effects were then used to determine which cognitive domains most strongly distinguish between males and females. In this way, our results can be directly related to findings from classical group studies. The highly predictive regions identified by the classification analysis were functionally characterised using the ‘Behavioural Domain (BD)’ categories available in the BrainMap database (http://brainmap.org/scribe/). Behavioural domains comprise main categories cognition, action, perception, emotion, and interoception, as well as their related sub-categories (Fox PT *et al.* 2014). Forward and reverse inference approaches were employed to determine the functional profile of the parcels with high classification accuracies. While forward inference is defined as the probability of observing activity in a brain region given knowledge of the psychological process, reverse inference is the probability of a psychological process being present given knowledge of activation in a particular brain region.

## 5 Results

### 5.1 Sex Prediction

#### 5.1.1 Within sample cross validation

For sample 1, across all parcels in the brain, the highest prediction accuracy reached 75.1%. All except five parcels’ accuracies were significant at p < 0.05 (FWE corrected for multiple comparisons) with a minimum accuracy of 63.1% and a mean prediction accuracy of 68.7% (S.D. 2.6%). The five non-significant parcels (accuracies between 61.5% and 62.9%) were located in right middle occipital gyrus, bilateral precuneus and right postcentral gyrus. The spatial distribution of classification accuracies across the whole brain is depicted in Figure 1 (a).

**Figure 1:**
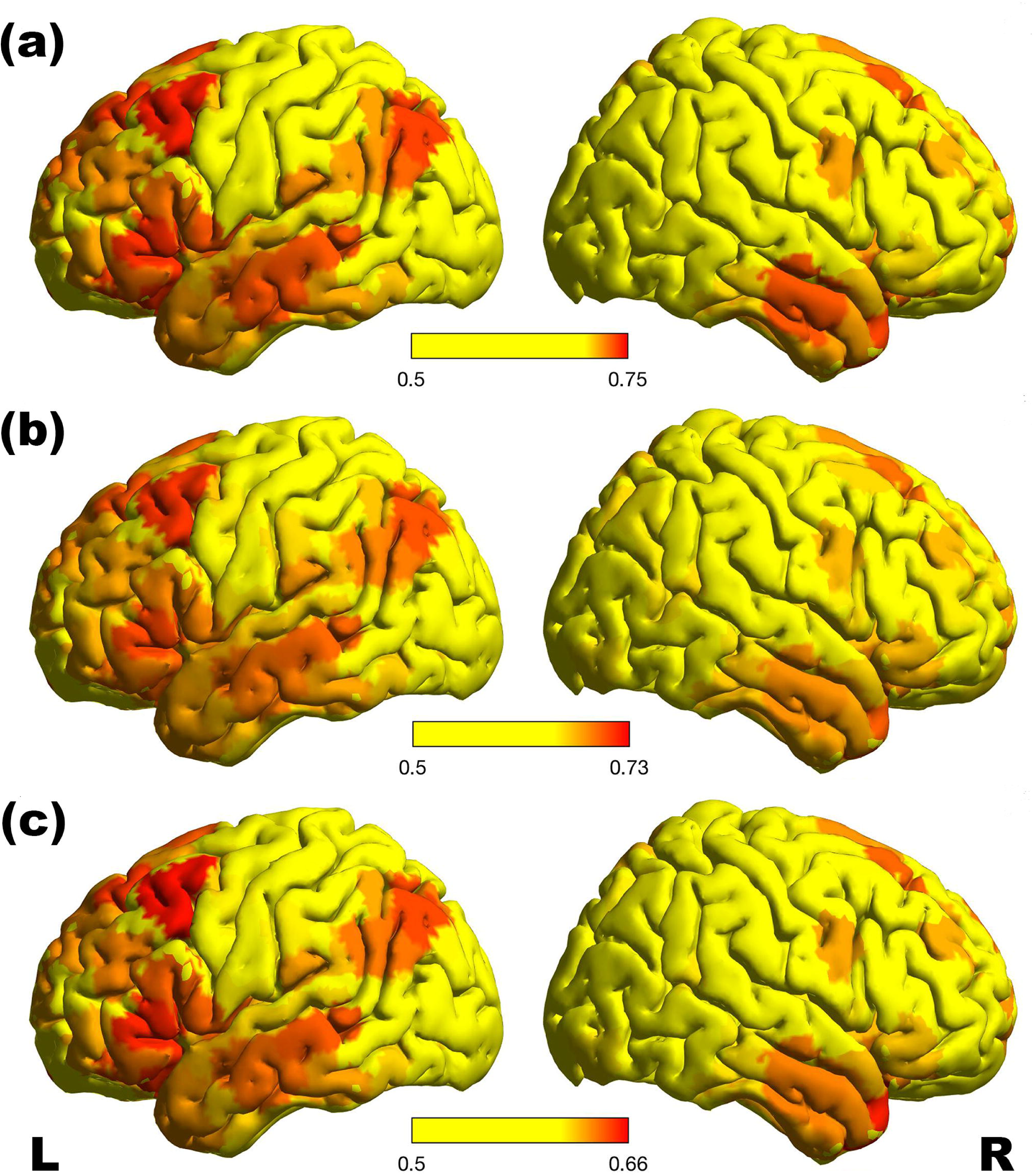
ROI based classification accuracies for within sample cross validation in sample 1 (a) as well as for across sample classification with the model trained on sample 1 and tested on sample 2 (b) and sample 3 (c). Across all analyses, the majority of the highly predictive parcels were located along the cingulate cortex, in right anterior mid-cingulate cortex as well as left posterior cingulate cortex. Other highly predictive parcels were located in bilateral medial frontal cortex and in bilateral precuneus. Further parcels with high prediction accuracy were located in left lateral frontal cortex, as well as left temporo-parietal regions and insula. The spatial distribution of parcels with highest classification accuracies were very similar across the within sample cross validation and the out-of-sample prediction for both samples, indicating the stability of the classification across samples with different characteristics.

#### 5.1.2 Between sample validation for sample 2

Across all parcels in the brain, the highest prediction accuracy reached 72.6%, with a minimum accuracy of 55.4% and a mean prediction accuracy of 64.3% (S.D. 3.0%). The spatial distribution of classification accuracies across the whole brain is depicted in Figure 1 (b).

#### 5.1.3 Between sample validation for sample 3

Across all parcels in the brain, the highest prediction accuracy reached 65.7%., with a minimum accuracy of 53.4% and a mean prediction accuracy of 60.0% (S.D. 2.5%). The spatial distribution of classification accuracies across the whole brain is depicted in Figure 1 (c).

Table 1 lists all those parcels, for which classification accuracy fell within the top 3% in all three analyses. The table lists the localization of all cortical regions that cover more than 10% of the respective parcel. Notably, the parcels with the highest classification accuracies were very similar across the within sample cross validation and the out-of-sample prediction for both samples. Consistency in the spatial distribution of highest classification accuracies in across sample validation can be taken to indicate the stability of the classification across samples with different characteristics. The stability of the spatial pattern of highly predictive parcels was further assessed by computing the rank correlation of within sample CV accuracy in sample 1 and between sample classification accuracy in samples 2 and 3 respectively. For both samples, correlation was highly significant (sample 2: r_s_ = 0.99, p > 0.0001; sample 3: r_s_ = 0.99; p< 0.0001). The scatterplots are shown in Figure 2.

**Figure 2:**
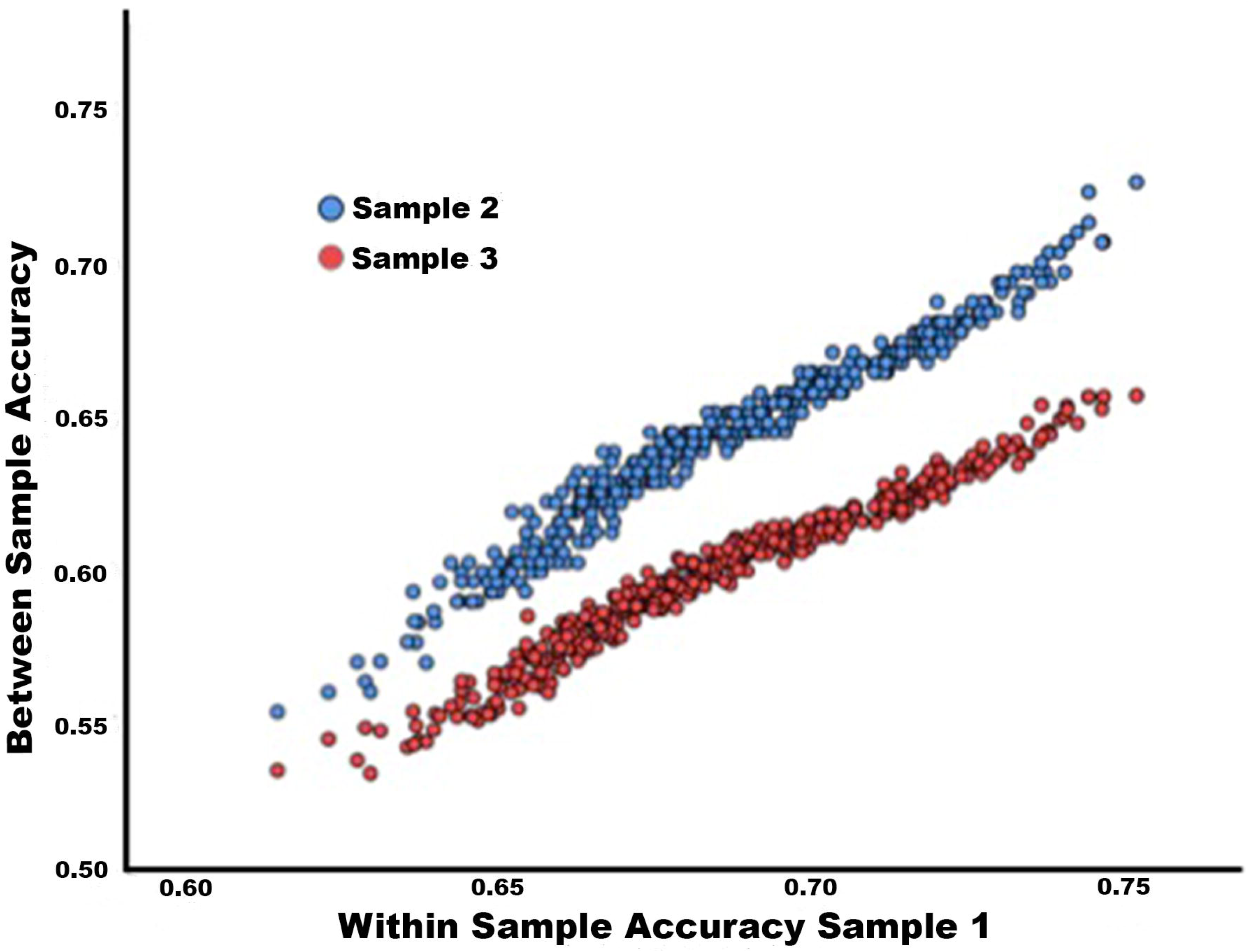
Scatter plot of classification accuracies for CV within sample one (HCP) versus out of sample classification in sample 2 (HCP, blue) and sample 3 (1000BRAINS, red) across the 436 parcels covering the whole brain.

**Table 1:** ROI based classification accuracies for brain regions with highest classification accuracies across analyses and significance values for forward and backward inference of the associated behavioral domains. Only domains significant at p < 0.05 (FDR-corrected) are given.

| Localization | Parcel<br>Size<br>(voxels) | Accuracy<br>(within<br>sample 1) | Accuracy<br>(sample1 -><br>sample 2) | Accuracy<br>(sample1 -><br>sample 3) | Mean<br>accuracy | Parcel<br>Nr. | Domains | P(Activation<br>/ Domain)<br>Likelihood<br>Ratio | P(Domain/<br>Activation)<br>Probability |
| --- | --- | --- | --- | --- | --- | --- | --- | --- | --- |
| R MCC (67.4%)<br>R ACC (19.4%) | 516 | 75.1% | 72.6% | 65.7% | 71.1% | 312 | Perception.Somasthesis.Pain<br>Action.Inhibito | 2.19<br>2.00 | 0.0466<br>0.0402 |
| R Precuneus (67.2%)<br>R MCC (29.4%) | 241 | 74.4% | 72.3% | 65.7% | 70.8% | 363 | Cognition.Social Cognition<br>Cogniton.Memory.Explicit<br>Emotion.Other | 3.34<br>1.95<br>1.93 | 0.764<br>0.1027<br>0.0933 |
| L Middle Frontal Gyrus<br>(99.0)% | 397 | 74.4% | 71.3% | 65.6% | 70.4% | 181 | Cognition.Memory.Working | 1.85 | 0.0856 |
| L Precuneus (67.8%)<br>L PCC (29.7%) | 205 | 74.6% | 70.7% | 65.7% | 70.3% | 154 | Cognition.Social Cognition<br>Cognition.Language | 4.10<br>3.60 | 0.0947<br>0.0194 |
|  |  |  |  |  |  |  | Emotion.Other | 2.36 | 0.1149 |
|  |  |  |  |  |  |  | Cognition.Memory.Explicit | 2.33 | 0.1240 |
| R Mid Orbital Gyrus<br>(64.3%)<br>R Rectal Gyrus (13.3%)<br>R ACC (10.5%) | 600 | 74.6% | 70.7% | 65.3% | 70.2% | 368 | Interoception.Thirst | 6.97 | 0.0104 |
|  |  |  |  |  |  |  | Perception.Gustation | 4.14 | 0.0453 |
|  |  |  |  |  |  |  | Emotion.Fear | 3.85 | 0.0398 |
|  |  |  |  |  |  |  | Cognition.Social Cognition | 2.86 | 0.0634 |
|  |  |  |  |  |  |  | Emotion.Reward | 2.64 | 0.1373 |
|  |  |  |  |  |  |  | Cognition.Reasoning | 1.80 | 0.1169 |
| L Mid Orbital Gyrus<br>(71.2%)<br>L Rectal Gyrus (23.2%) | 344 | 74.0% | 70.7% | 65.4% | 70.0% | 161 | Emotion.Reward | 2.87 | 0.1511 |
|  |  |  |  |  |  |  | Cognition.Social Cogniton | 2.44 | 0.0548 |
| R ACC (70.6%)<br>R Superior Medial<br>Gyrus (13.5%) | 435 | 74.2% | 71.0% | 64.8% | 70.0% | 370 | Perception.Gustation | 4.15 | 0.0470 |
|  |  |  |  |  |  |  | Emotion.Reward | 2.33 | 0.1256 |
|  |  |  |  |  |  |  | Cognition.Social Cogniton | 2.16 | 0.0497 |
| L IFG p. Orbitalis<br>(19.8%)<br>L IFG p. Triangularis | 343 | 74.0% | 70.7% | 65.2% | 70.0% | 186 | Cognition.Language.Syntax | 3.35 | 0.0275 |
|  |  |  |  |  |  |  | Cognition.Language | 2.75 | 0.0154 |
|  |  |  |  |  |  |  | Cognition.Language.Semantics | 2.57 | 0.1764 |
| (77.6%) |  |  |  |  |  |  | Cognition.Language.Speech | 1.91 | 0.0867 |
|  |  |  |  |  |  |  | Cognition.Memory.Explicit | 1.49 | 0.0826 |
| L Superior Frontal<br>Gyrus (90.2%) | 246 | 73.9% | 70.4% | 64.9% | 69.7% | 152 | Cognition.Social Cognition | 3.34 | 0.0756 |
|  |  |  |  |  |  |  | Action.Inhibition | 2.73 | 0.0496 |
| L IFG (p. Orbitalis<br>(67.9%)<br>L Insula Lobe (19.0%)<br>L Temporal Pole<br>(12.3%) | 358 | 74.0% | 69.7% | 65.0% | 69.6% | 183 | Emotion.Disgust | 2.82 | 0.0180 |
|  |  |  |  |  |  |  | Cognition.Social Cognition | 1.85 | 0.0448 |
|  |  |  |  |  |  |  | Cognition.Language.Semantics | 1.69 | 0.1170 |
| L Angular Gyrus<br>(60.9%)<br>L Inferior Parietal<br>Lobule (20.9%)<br>L Middle Occipital<br>Gyrus (12.5%) | 297 | 73.7% | 70.4% | 64.5% | 69.5% | 150 | Cognition.Memory.Explicit | 1.89 | 0.1046 |

Furthermore, to examine the consistency of the classification patterns across samples, we computed the mean parcelwise connectivity pattern for females and males respectively within each of the three samples. By subtracting female mean connectivity from male mean connectivity per parcel, the mean sex difference in connectivity was computed as a 1×x 435 (number of parcels – 1) vector for each of the three samples. Then the sex difference in connectivity for sample 1 was correlated with those for sample 2 and sample 3 respectively. For both samples, the correlation was significant, indicating comparable patterns of connectivity between parcels (sample 1 / sample 2: r = 0.601, p < 0.001; sample 1 / sample 3: r = 0.259, p < 0.001). The difference in correlation strengths between sample 2 und sample 3 presumably reflects the fact that sample 1 and sample 2 were both drawn from the same samples, while sample 3 constituted a fully independent sample with different participant and scanning characteristics.

### 5.2 Prediction accuracies for the whole brain connectome

To compare the parcelwise classification performance to accuracies that can be achieved based on the whole brain connectome, respective analyses were run on the full connectome derived from the 436 parcels. While within-sample cross-validation within sample 1 achieved an accuracy of 74.81%, the between sample validation showed an accuracy of 70.39% for sample 2 as well as of 51.13% for sample 3.

### 5.3 Confounding effects of grey matter volume

To examine the influence of volumetric differences between the sexes, a standard analysis pipeline of the CAT12 toolbox (http://www.neuro.uni-jena.de/cat/) was used to compute GMV for each parcel in each participant within sample 1, based on their T1-weighted anatomical scan (3D MPRAGE, TR = 2400ms, TE = 2.14ms, FOV 224mm × 244mm, voxel size = 0.7 mm isotropic). This data was used to compute the mean sex difference in GMV in each parcel by subtracting, for each parcel, mean GMV across females from mean GMV across males. Then, a rank correlation was computed between the mean sex difference in GMV and RS-based classification accuracy across parcels. This was done for both the directed as well as the absolute mean sex difference in GMV. Both correlations were non-significant (directed sex difference: r =0.0374, p =0.4364; absolute sex differences: r = −0.0320, p = 0). Similar correlations were computed across those parcels for which classification accuracy was significant. Again, both for directed and absolute sex differences these correlations were non-significant (directed sex difference: r = 0.0264; p = 0.5851; absolute sex difference: r = −0.0210, p = 0.6641). Finally, the correlations were computed across the parcels with top 10% classification accuracies, again not revealing any significant correlations (directed sex difference: r = 0.0909; p = 0.5561; absolute sex difference: r = −0.0519, p = 0.7374).

Furthermore, for each parcel, an independent samples t-test was used to compare GMV for those subjects which were correctly classified to those which were misclassified. While 21 of the comparisons were significant at p < 0.05, none remained significant after FDR correction for multiple tests across the 436 parcel, indicating that parcel-wise GMV was not significantly different between correctly and incorrectly classified subjects.

Additionally, we repeated the cross-validation analysis on sample 4, in which groups were matched for grey matter volume. Overall, classification accuracies in sample 4 were lower than in sample 1. Considering that sample 4 was constructed from subsets of sample 1 and sample 2, similar SNR can be expected in these samples. Thus, it is assumed that that lower accuracies observed for sample 4 are based on the smaller sample size, as it has been shown that, when a sample is fixed then smaller random sub-samples will show a lower accuracy as there is simply less data to learn from (Provost K et al. 1999). Therefore, while smaller sample sizes tend to lead to higher variance in accuracies, they also, on average, result in lower accuracies. Even though, sex classification could still be achieved across the whole brain with accuracies between 55.6% and 71.8% (mean accuracy 64.7%). Furthermore, the overall pattern of parcels with highest classification accuracies was comparable to our original results as shown in Supplementary Figure 1.

### 5.4 Functional Decoding

Functional characterization according to the BrainMap meta-data was performed for all parcels that appear within the top 3% of parcel accuracies in the within sample CV as well as the between sample classification.

#### 5.4.1 Anterior cingulate and medial frontal areas

Parcels that covered regions in the right medial frontal cortex as well as anterior and middle cingulate cortex were mainly associated with behavioural domains (BDs) emotion, specifically fear and reward, cognition, especially social cognition and reasoning, perception, including gustation and pain and interoception, especially thirst, as well as action inhibition.

#### 5.4.2 Middle and posterior cingulate and precuneus

The parcel in the right middle cingulate gyrus and precuneus was associated with BDs of emotion, as well as social cognition and explicit memory. Similar BDs were associated with the parcel in the left posterior cingulate cortex and precuneus.

#### 5.4.3 Left lateral frontal areas

Parcels in the left lateral frontal cortex were associated with BDs of working and explicit memory, as well as speech and language (especially semantics), social cognition and emotion (disgust) as well as action inhibition.

#### 5.4.4 Left angular gyrus

The parcel centred in the left angular gyrus was associated with BDs of explicit memory.

In summary, those parcels for which connectivity patterns with the rest of the brain achieved highest sex classification accuracies were associated with different types of emotion, social cognition, memory and language. The BDs associated with the top accuracy parcels are summarized in **Figure 3.**

**Figure 3:**
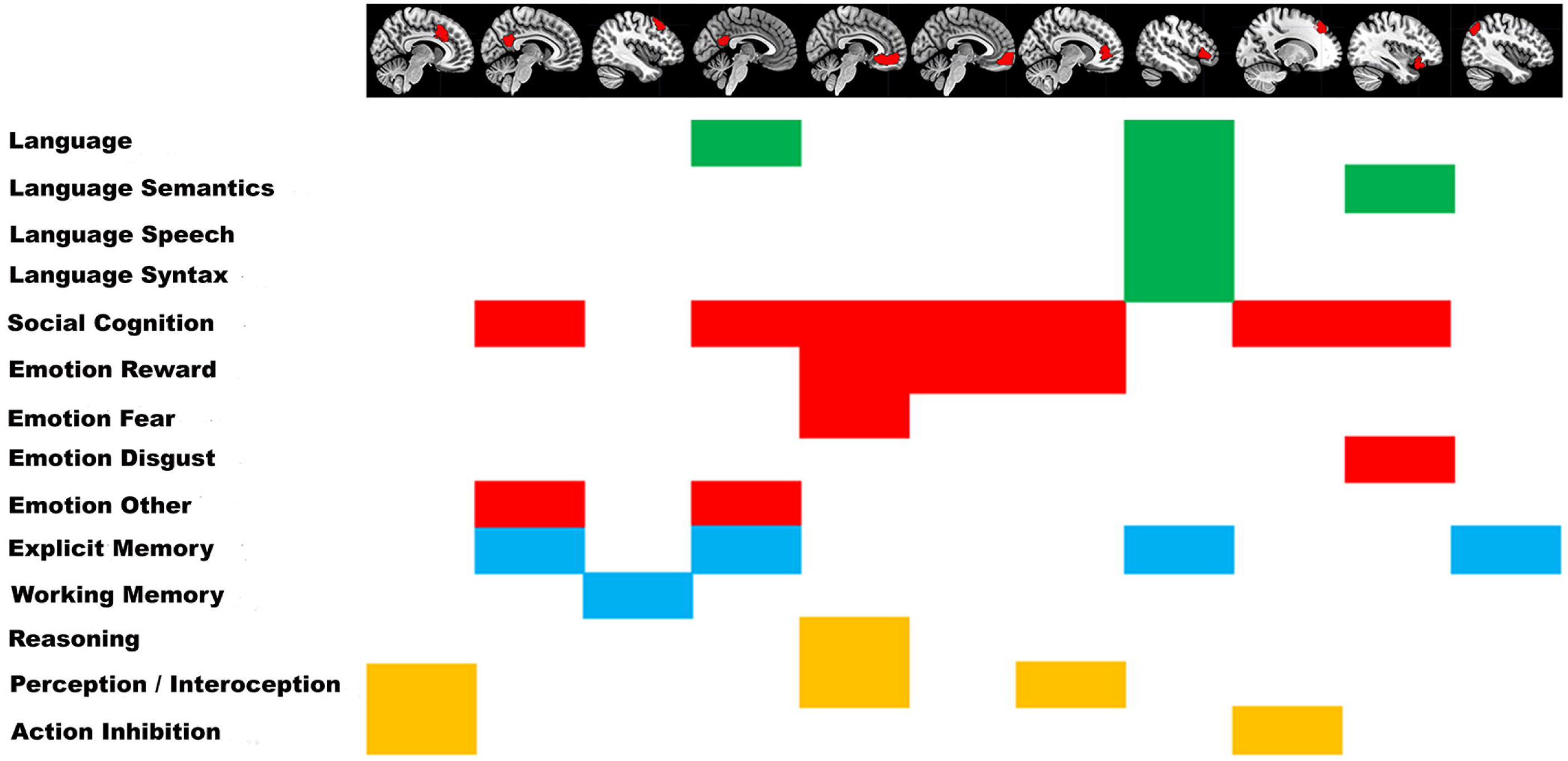
Behavioural domains associated with the brain parcels that achieved highest sex classification accuracies, as defined by functional decoding.

## 6 Discussion

Our data revealed that classification of an out-of-sample subjects’ sex from the RS connectivity profiles was possible with high accuracies across the whole brain, indicating that each individual parcel’s RS connectivity carries enough information to reliably identify a previously unseen subject’s sex. However, and most importantly, there were pronounced differences in the prediction accuracies of specific brain regions, indicating spatially specific effects. Of note, those spatially specific parcels with high prediction accuracies were stable across within and between samples classification, indicating the generalizability of these predictions independently of the specific characteristics of the sample as well as the specific imaging parameters used. These results strongly indicate, that it is the functional connectivity of specific regions in the brain that is most characteristically different between males and females.

We employed a functional decoding approach to examine which cognitive domains were related to those brain regions that most clearly differentiate between the sexes. As opposed to group studies comparing brain activations and cognitive performances, our approach does not rest on the assumption of a clear-cut sexual dimorphism in functional brain organization, but is suitable to characterize multi-layered differences in functional connectivity of certain brain regions, while making it possible to assess cognitive domains in which males and females differ most.

Of note, functional connectivity in itself is obviously not spatially specific. Thus, high classification accuracy for a given parcel does not result from the intrinsic function of that parcel, but rather from its pattern of connectivity with other parcels across the brain. However, there is general consensus that mental functions arise from the coordinated activity within distributed networks rather than any individual brain region (Park HJ and K Friston 2013). A high classification accuracy for a given parcel indicates differences in this parcel’s connectivity, which in turn can be taken as evidence that those cognitive domains, for which the parcel is typically activated, is organized in different ways in males than in females or at the very least that the pattern of that cognitive function is more “typical” (i.e. less variable) in one of the sexes as opposed to the other.

Similarly, as the present analyses are based on connectivity patterns across the whole brain, there is a large amount of shared information between parcels, due to the fact that connectivity with other brain parcels is used as features in the analysis. This dependency between parcels presumably is reflected in a relatively uniform classification performance across the brain with only very few parcels displaying non-significant discrimination. Still, our parcelwise classification approach clearly identified a subset of parcels that most strongly distinguished between males’ and females’ brain connectivity patterns and thus allowed for an identification of cognitive domains which can be assumed to most strongly differentiate between the sexes.

### 6.1 Spatially specific effects

Across all analyses, the majority of the highly predictive parcels were located along the cingulate cortex, in right anterior mid-cingulate cortex as well as left posterior cingulate cortex. Other highly predictive parcels were located in bilateral medial frontal cortex, in bilateral precuneus as well as left lateral frontal cortex, left temporo-parietal regions and insula.

Interestingly, the majority of parcels showing highest classification accuracies in the present study match well with brain areas that have been related to the default mode network (Buckner RL et al. 2008; Biswal BB *et al.* 2010). One of the largest studies conducted on the topic (Biswal BB et al. 2010) indicated that women exhibited stronger connectivity than men in the posterior cingulate cortex, medial prefrontal cortex and the inferior parietal lobe, but weaker connectivity in the dorsal anterior cingulate cortex, insula, superior temporal gyrus, superior marginal gyrus and occipital regions. Our results are in line with these findings (Biswal BB et al. 2010), as most of the regions reported in their study exhibit high classification accuracies here. Furthermore, given that the methods employed in both studies to assess functional connectivity patterns in the RS are different, the overlap of results further speaks to the generalizability of our findings.

Additionally, the importance of the DMN in the classification of participant’s sex based on RS fMRI data replicates findings from (Zhang C *et al.* 2018). These authors employed whole brain connectivity in their classification approach and subsequently identified those connections, which contributed most to the successful classification. While our approach is different in that it is based on spatially specific connectivity, both studies identified the importance of the DMN in successful sex classification. The present results also match with findings from a large-scale group comparison based study (Ritchie SJ *et al.* 2018) which showed that connectivity within the DMN is more pronounced in females than in males.

Further brain regions which play an important role in successful sex classification are associated with cognitive domains for which sex differences have previously been identified. For examples, brain regions displaying high accuracies in differentiating between males and females in the present study closely match with those reported in classical studies suggesting language processing as the key cognitive domain to differ between males and females (Halpern D 1992). For example, previously reported brain regions which were also identified in the present study include the angular gyrus, the prefrontal and retrosplenial cortex (Frost JA et al. 1999), as well as the (pre-)cuneus and cingulate areas (Clements AM et al. 2006). Thus, our results support the existence of differences in the brain basis for language processing as one of the most distinguishing features between the sexes.

Further high prediction accuracies were identified for medial brain regions, specifically in the frontal cortex. In accordance with our results, a meta-analysis of the neural correlates of sex differences in emotion processing (Stevens JS and S Hamann 2012) identified several of the brain regions which provided highly accurate sex prediction in our study, like the medial frontal and anterior cingulate regions.

Finally, the high classification accuracy in bilateral precuneus might be linked to the established male advantage in visuo-spatial working memory, which has recently been demonstrated based on a large meta-analysis (Voyer D et al. 2017).

Altogether, similar to (Zhang C *et al.* 2018), our findings show that accurate sex prediction is possible on the basis of brain connectivity at rest. However, in addition to their findings, our novel approach enabled identification of brain regions, for which the connectivity with the rest of the brain is most distinctive between males and females. Functional decoding of these regions identified the cognitive domains associated with these regions. Speaking to the reliability of our findings, these domains match well with those, for which sex differences have previously been reported based on group comparison. However, importantly, our results emphasize the importance of these areas for sex differences in a much more direct way.

However, it needs to be noted that spatial differences in the signal-to-noise ratio (SNR) of the fMRI data might influence the results. For example, for regions with low SNR we might not be able to achieve good accuracies, even if biologically speaking, they are important for distinguishing male versus female. Future studies might want to take into account measures of SNR across the brain. While this problem is not specific to the approach employed here but rather exists for any analysis of fMRI data, it underlines the importance of a thorough quality control of the data, specifically for these types of studies.

### 6.2 Accuracy and Generalizability of sex classification

Firstly, our samples specifically excluded pairs of related subjects. We could therefore make sure that classification accuracies are not optimistically biased because subjects are related. Due to this selection, the sample on which our model was trained is much smaller than the sample employed in (Zhang C *et al.* 2018), in fact is contains only about half as many subjects. Furthermore, we decided to base our predictions on the first of the available RS runs only. While for the HCP data four runs of RS data are available, this is not the case for most other data sets, to which this method might be applied. We aimed to not mainly identify the maximum accuracy that can be achieved – for example by combining several RS runs as done by (Zhang C *et al.* 2018) did, but rather to show that successful classification is possible based on just one run of RS data. Indeed, our results show, that high classification accuracies can be achieved based on relatively small samples and just about 10 minutes of RS data.

More importantly, we could show that the ability of spatially specific brain regions to predict sex is stable not only within sample but also across different samples. The HCP sample and the 1000BRAINS sample differ with respect to both imaging parameters and sample characteristics. For example, while the HCP sample contains only relatively young participants, the 1000BRAINS sample comprised a much wider age range including older subjects. Still, the highest within-sample and between-sample accuracies were found in highly similar brain regions, underlining the reliability of our findings. Furthermore, average sex differences in spatially specific brain connectivity patterns appear to be comparable across sample, indicating that not only the capacity to classify, but also the underlying connectivity patterns show similarities across samples with different characteristics. Still, with the availability of more large data set, it would further strengthen the results if other groups could independently replicate the same spatial patterns.

It needs to be noted that total brain volume is one variable demonstrating a consistent sex difference and thus might have influenced prediction accuracies as observed here. This issue has been previously addressed by (Zhang C et al. 2018), who showed that both sex and brain volumes could be predicted from resting state brain connectivity across the whole brain. Based on showing that features (i.e. functional brain connections) in sex and brain volume predictions overlap by less than 20%, these authors concluded that the sex difference in brain volume is not dominating in gender sex prediction. However, this analysis does not address spatially specific sex differences in GMV, which might have influenced the parcelwise classifications examined here. While spatially specific sex differences exist in the data used in the present analyses, these local differences in GMV are unrelated to parcelwise classification accuracies, indicating that the quality of the classification is independent of local volumetric differences between the sexes. There was also no systematic association between parcelwise GMV and individual classification performance, further suggesting that local GMV did not influence our results.

Finally, a classification analysis in a sample that was matched for grey matter volumes between males and females, displayed slightly lower accuracies, but a similar spatial pattern of highly predictive parcels as the original analysis, again indicating that grey matter volume did not influence the classification.

### 6.3 Does a sexual dimorphism of functional brain organization exist?

Our results show that accurate prediction of the sex of an out-of-sample subject is possible based on individual brain regions’ RS connectivity. The spatially specific effects identified here are closely linked to sex differences in cognition.

Our data show that classification based on specific brain regions can achieve classification accuracies that are comparable or even higher than what can be achieved based on the whole brain connectome. What is more, when considering the whole brain classification accuracies, the drop in accuracy between within-sample cross validation and across sample performance (especially in the independent sample) is more pronounced than for the parcelwise analysis, which might indicate an overfitting based on the extremely high dimensionality of the whole brain connectome.

While the vast majority of parcels distinguish between males and females with significant accuracy, for none of the parcels’ prediction accuracies were approaching 100%. One reason for the non-perfect prediction accuracies might be based on the fact that our approach ignores functional brain networks. Thus, while we cannot exclude that this might be based on methodological choices with respect to the machine learning approach, it might also add further support to the recent suggestion (Joel D *et al.* 2015), that, even for specific regions, brains falling on the ends of the male-female continuum are rather rare. While we cannot directly test this assumption based on the present data sets, it is conceivable that where a brain falls on this continuum, might be modulated by effects of each individual’s experience, education, and culture or a combination of these (Jancke L 2018).

Thus, our results do not support an actual sexual dimorphism of the human brain. Same as for brain structure (Joel D *et al.* 2015), features based on RS connectivity appear to substantially overlap between males and females. Specifically, our data indicates, that while some regions of the brain distinguish better between the male and the female brain, it appears to be impossible to actually identify dimorphic features that justify a clear sex distinction. In fact, this is not surprising but rather might suggest that the functional organization of each individual brain is related to the individual’s sex, but is also shaped by additional factors. Further research needs to elucidate the biological and social factors contributing to each individual’s specific brain organization pattern.

For example, one of the most obvious biological factors influencing sex differences in functional brain organization might be hormonal differences, for example fluctuating sex hormones across the menstrual cycle in women. In fact, a variety of studies have shown hormonal effects on functional brain connectivity RS fMRI studies (Hjelmervik H et al. 2012; Petersen N et al. 2014; Arelin K et al. 2015; De Bondt T et al. 2015; Weis S et al. 2017). With locally specific functional brain organization varying with hormonal chances, females might be expected to exhibit increased variability in those regions that contribute most to sex classification. This in turn might have a profound influence on classification performance, with classification accuracies possibly depending on the female participants’ cycle phase. It might also mean that successful classification can only be achieved in specific cycle phases. This is a limitation of the present study, and in fact any study assessing sex differences without consideration of the females’ menstrual cycle. Unfortunately, so far none of the large-scale data sets necessary for the analyses like those presented here, has collected information on hormone levels. If this could be done in the future, hormonal information might in fact further inform the models and thus increase classification accuracies. Furthermore, it would be highly interesting to examine if hormonal changes are region specific or affect classification performances across the whole brain.

Also, there might be social factors influencing classification accuracies and regionally specific effects, such as individual learning experiences, culture, or gender stereotypes. In fact, it might be the individual pattern of interaction between biological and social factors that is picked up by the classifier. Based on our data alone, it is not possible to disentangle these modulating effects. Thus, future studies, should take into account not only the biological sex and biological modulators like genetics and hormones, but also social factors like the self-perceived gender of the participants. Presumably, only the combination and interaction of all these factors will enable a more detailed characterization of individual variations in functional brain organization.

### 6.4 Conclusions

Our results show that sex classification based on RS fMRI data is possible with high accuracies, which are significantly different from chance across more or less the whole brain. The results also show that classification can be reliably extended to independent samples, differing both with respect to imaging parameters and sample characteristics. Those regions that display high prediction accuracies are stable across samples, indicating that the spatial pattern of regions that best distinguish males from females generalizes across samples and age-ranges. This is further underlined by the fact that these regions confirm sex differences that have been shown in classical group comparison. In addition, they match well with areas that have been related to specific clinical conditions for which prevalence differs between the sexes.

However, our results also indicate that sex alone cannot perfectly explain each individual’s specific patterns of functional brain organization. Thus, these data do not support the existence of a sexual dimorphism with respect to functional brain organization and they strongly support the notion that terms such as “female brains” or “male brains”, which are frequently used especially in popular writing, are not appropriate. While some patterns of brain organization might be driven by sex, more complex pattern of brain organization are most likely shaped by each individual’s environment and experiences and thus cannot be explained by sex alone.

## 7 Acknowledgments

This study was supported by

1. the Deutsche Forschungsgemeinschaft (DFG, EI 816/11-1),
2. the National Institute of Mental Health (R01-MH074457),
3. the Helmholtz Portfolio Theme “Supercomputing and Modeling for the Human Brain”,
4. the European Union’s Horizon 2020 Research and Innovation Programme under Grant Agreement No. 720270 (HBP SGA1) 785907 (HBP SGA2).
5. BTTY is supported by Singapore National Research Foundation (NRF) Fellowship (Class of 2017).

---

**Supplementary Figure 1:** ROI based classification accuracies for within sample cross validation in sample 4. While the classification accuracies are lower than for sample 1, the spatial distribution is comparable to the within-sample cross validation in sample 1.

