## Supplementary figures and images for "Sex classification by resting state brain connectivity"

### Supplementary Figure 1

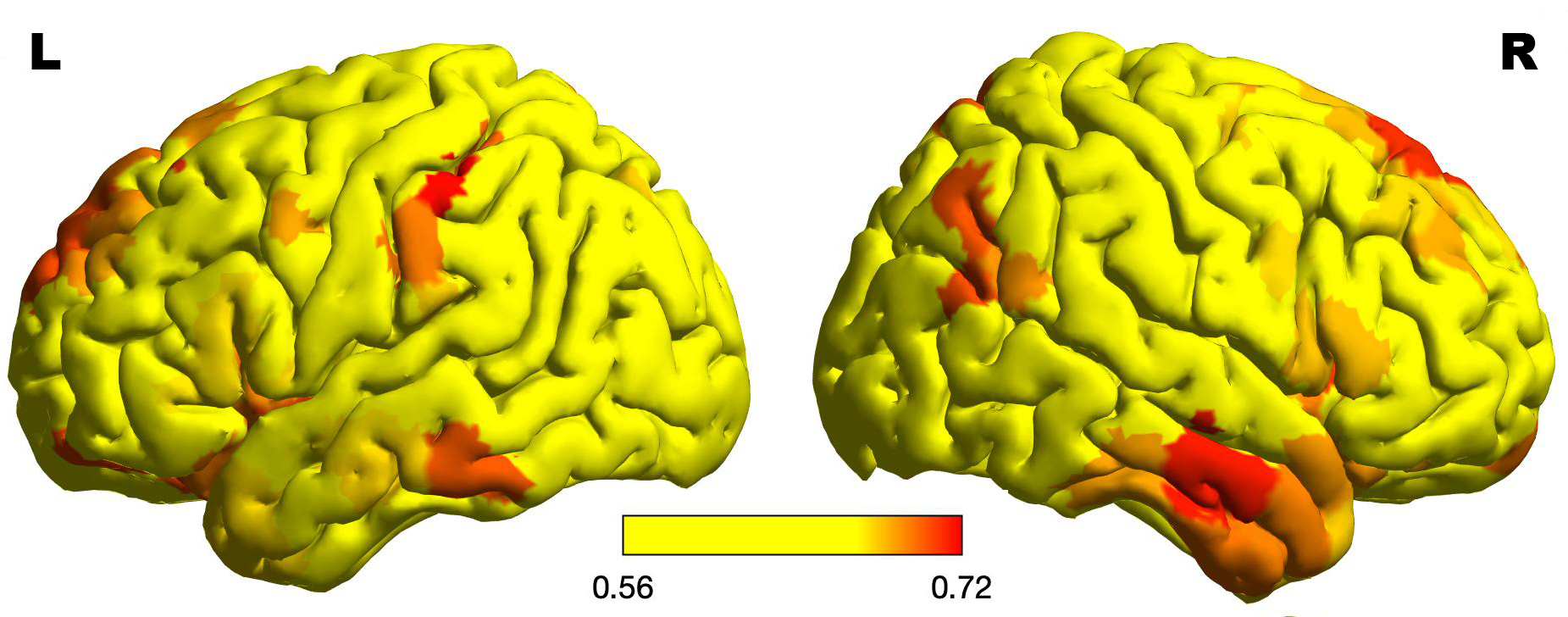
